# Single-cell analyses of immune thrombocytopenic patients reveal multiorgan dissemination of high-affinity autoreactive plasma cells

**DOI:** 10.1101/2021.06.29.450355

**Authors:** Pablo Canales-Herrerias, Etienne Crickx, Matteo Broketa, Aurélien Sokal, Guilhem Chenon, Imane Azzaoui, Alexis Vandenberghe, Angga Perima, Bruno Iannascoli, Odile Richard-Le Goff, Carlos Castrillon, Guillaume Mottet, Delphine Sterlin, Ailsa Robbins, Marc Michel, Patrick England, Gael A. Millot, Klaus Eyer, Jean Baudry, Matthieu Mahevas, Pierre Bruhns

## Abstract

The major therapeutic goal for immune thrombocytopenia (ITP) is to restore normal platelet counts using drugs to promote platelet production or by interfering with mechanisms responsible for platelet destruction. 80% of patients possess anti-integrin αIIbβ3 (GPIIbIIIa) IgG autoantibodies causing platelet opsonization and phagocytosis. The spleen is considered the primary site of autoantibody production by autoreactive B cells and platelet destruction. The immediate failure in ~50% of patients to recover a normal platelet count after anti-CD20 Rituximab-mediated B cell depletion and splenectomy suggest that autoreactive, rituximab-resistant, IgG-secreting B cells (IgG-SC) reside in other anatomical compartments. We analyzed >3,300 single IgG-SC from spleen, bone marrow and/or blood of 27 patients with ITP revealing high inter-individual variability in affinity for GPIIbIIIa with variations over 3 logs. IgG-SC dissemination and range of affinities were however similar per patient. Longitudinal analysis of autoreactive IgG-SC upon treatment with anti-CD38 mAb daratumumab demonstrated variable outcomes, from complete remission to failure with persistence of high-affinity anti-GPIIbIIIa IgG-SC in the bone marrow. This study demonstrates the existence and dissemination of high-affinity autoreactive plasma cells in multiple anatomical compartments of patients with ITP that may cause the failure of current therapies.

## Introduction

Immune Thrombocytopenia (ITP) is an immune disorder characterized by a strong autoimmune response against platelet autoantigens that causes platelet destruction^1^. Clinical manifestations often present as mild bleeding on the skin or mucosal surfaces, but can also include life-threatening internal bleeding episodes in more severe cases^2,3^. The hallmark of ITP is the presence of anti-platelet antibodies, which contributes to the accelerated destruction of platelets by mechanisms including FcR-mediated phagocytosis by macrophages, and inhibition of new platelet generation by megakaryocytes. Autoantibodies against platelets are predominantly of the IgG isotype^4^, with IgG1 being the most prevalent subclass^5^. Although several platelet surface proteins are known to be targeted by autoantibodies, namely GPIb/IX/V, GPIa/IIa and GPIV^6^, the glycoprotein GPIIbIIIa has long been known as the dominant autoantigen in ITP, with anti-GPIIbIIIa antibodies found in 60-90 % of patients^6–8^.

The spleen plays a major role in the pathophysiology of ITP^9^ and splenectomy remains the most effective and curative therapy^10^. Indeed, autoreactive anti-GPIIbIIIa-secreting cells arise from germinal center reactions located in the spleen of patients with ITP. Germinal center expansion results from T follicular helper cell activation^11^ and reduced T regulatory cell numbers^12^. Pathogenic anti-GPIIbIIIa antibody-secreting cells can be detected by ELISPOT in the spleen and blood of patients with ITP^9,13,14^. In this prototypic antibody-mediated autoimmune disease, B cell depletion using rituximab (anti-CD20 mAb) is widely used with up to 60% of patients having a short-term complete response^15^, 40% achieving durable responses (6 – 12 months) and only 20 to 30% achieving a long-term response (5 years)^16,17^. Paradoxically, B cell-depletion therapy stimulates an adaptation of splenic short-lived plasma cells leading to their reprogramming into long-lived spleen cells in patients with ITP, explaining in part primary failure of rituximab^13^. Furthermore, relapse of ITP after B cell depletion therapy by rituximab usually occurs during B cell lymphopoiesis (more than 6 months after treatment), corresponding to the re-initiation of an autoimmune B cell response. We recently demonstrated that rituximab-resistant memory B cells directly contributed to ITP relapses by enabling autoreactive germinal centers along with the recruitment of naïve B cells and leading to anti-GPIIbIIIa secretion by newly formed antibody-secreting cells (ASC)^18^.

In a T-dependent response such as ITP, ASC arise from germinal centers or memory B-cells and egress out of spleen as short-lived ASC to circulate in the blood, with some maturing into long-lived plasma cells (LLPCs) in the bone marrow. The longevity of ASC is not primarily cell-intrinsic but largely depends on signals provided by their microenvironment^19–21^. Despite their fundamental role, a major impediment to studying heterogeneity of the autoimmune ASC is the difficulty to assess the specificity of such cells that express no or relatively few immunoglobulins at their surface. In order to decipher the exact contribution of different subsets of anti-GPIIbIIIa ASC to the pathogeny in different organs and in different clinical situations, we performed herein a high-throughput phenotypic analysis of these autoreactive human IgG-secreting cells (IgG-SC) at single-cell level, comparing patients with ITP and healthy donors. We describe a 3-log repertoire of affinities of anti-GPIIbIIIa IgG autoantibodies secreted from freshly isolated spleen, bone marrow and blood. The proportion of autoreactive cells among IgG-SC highly correlated between the spleen and the two other compartments, and very high-affinity anti-GPIIbIIIa IgG-SC were detected in all three compartments. A longitudinal analysis of autoreactive IgG-SC in 3 patients refractory to approved treatments who were administered anti-CD38 daratumumab demonstrated inter-individual variability in the targeting of high-affinity anti-GPIIbIIIa IgG-SC. These studies identify the large repertoire of anti-GPIIbIIIa affinities in patients with ITP and demonstrate the existence and dissemination of high-affinity autoreactive ASC to the bone marrow of patients with ITP that may be the underlying cause of failure of current therapies.

## Results

### Single-cell bioassay allows phenotypic characterization of autoreactive ASC

In order to characterize directly the secretion rate, specificity, and affinity for GPIIbIIIa of IgG secreted by autoreactive plasma cells and plasmablasts, collectively termed ASC hereunder, without the need to sort, clone, or re-express antibodies, we adapted a single-cell bioassay in microfluidic droplets (termed “DropMap”) that we have described previously^22,23^. Mononuclear cells from the spleen, bone marrow, or blood of patients and fluorescent bioassay reagents are co-encapsulated in droplets, immobilized within an observation chamber and imaged over 1 hour by time-lapse fluorescence imaging. Magnetic beads in each droplet align into a line (beadline) under a magnetic field to serve as a physical surface for a double-fluorescent sandwich ELISA revealing IgG secretion from the cell and specificity of that IgG for GPIIbIIIa (Figure 1a). The time-resolved fluorescence signals allow for estimation of IgG secretion rates and affinity for GPIIbIIIa of the secreted IgG by using calibration curves generated with monoclonal anti-GPIIbIIIa IgG of known affinity (K_D_). The anti-GPIIbIIIa reference curve, generated using nine anti-GPIIbIIIa IgG mAbs of various affinities, allows for measurements over 2 logs of affinities i.e., 2.5×10^−10^ ≤ K_D_ ≤ 5×10^−8^ M (Figure 1b). Therefore herein, IgG interacting with GPIIbIIIa at a calculated K_D_ below 5×10^−8^ M will be considered binding i.e., anti-GPIIbIIIa IgG antibodies, and the cell secreting such IgG will be termed GPIIbIIIa-reactive ASC.

**Figure 1.**
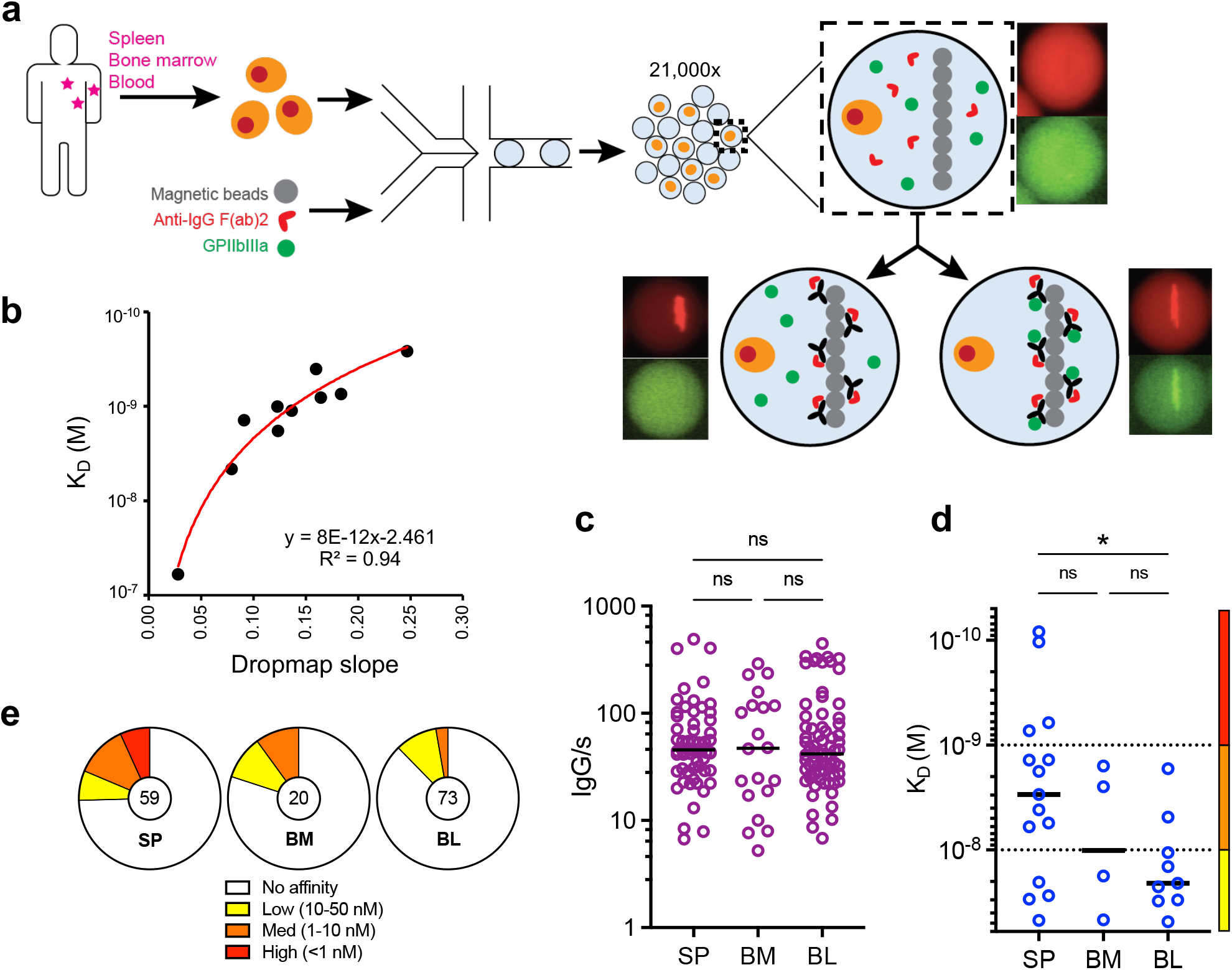
Single cell analyses reveal secretion rate and GPIIbIIIa affinity from IgG-SC. (**a**) Schematic of the DropMap pipeline. Single mononuclear cells are encapsulated in droplets, together with magnetic beads coated with anti-kappa light chain nanobody (VHH), fluorescently-labeled anti-IgG F(ab’)2-Alexa647 (red) and GPIIbIIIa-Alexa488 (green). Droplets are immobilized in the chip, exposed to a magnetic field to induce beads to form a beadline, and imaged over time. (**b**) GPIIbIIIa affinity reference curve generated using anti-GPIIbIIIa mAbs with known K_D_ and values obtained from DropMap experiments using these mAbs. (**c-e**) DropMap analysis of spleen, blood and bone marrow samples from patient F at the time of splenectomy: (**c**) IgG secretion rate; (**d**) affinity for GPIIbIIIa (K_D_) classified into “high” (red), “medium” (orange) and “low” (yellow) affinity with dotted lines separating these categories; (**e**) distribution of IgG-SC into low (yellow), medium (orange) and high (red) affinity binders to GPIIbIIIa or non-binders (white), with total IgG-SC numbers indicated. (**c-d)** Single cell values and medians are plotted. NS (not significant); *P < 0.05; **P < 0.01; ***P < 0.001 using a Welch test and multiple testing P value adjustment. See Supplemental Table 2 for further details.

As an example, samples from the spleen, bone marrow, and blood were obtained at the time of splenectomy from one patient (patient “F”; 73 y/o, male) who was splenectomized 8 months after last rituximab infusion due to treatment failure (Fig.1c-e). All data from these samples were acquired in duplicates on the same day with high reproducibility in total and autoreactive IgG-SC detection per replicate (Supplementary Figure S1). For each organ we analyzed 5,000-9,000 single cells in total, and found IgG-SC represented 0.55%, 0.41%, and 1.26% of the mononuclear cell pool in the spleen, bone marrow, and blood, respectively. IgG-SC from the three organs displayed a similar range (5-500) and median values (~50) of IgG secretion in molecules per second (IgG/s) (Figure 1c). Within the IgG-SC pool of all three organs, a fraction secreted anti-GPIIbIIIa IgG antibodies with K_D_ values in the 5×10^−8^ M – 1×10^−10^ M range that we categorized into low- (50nM ≤ K_D_ ≤ 10nM), medium- (10nM < K_D_ ≤ 1nM) and high-affinity (K_D_ < 1nM) GPIIbIIIa-reactive IgG-SC to facilitate subsequent analyses (Figure 1d). In this patient, the spleen contained IgG-SC with significantly higher affinities for GPIIbIIIa as compared to the blood, and the proportion of GPIIbIIIa-reactive cells among IgG secreting cells was 25% in the spleen, 20% in the bone marrow and 12% in the blood, with high-affinity antibodies detected only in the spleen (Fig 1e).

### High inter-individual variability of autoreactive ASC presence and affinity in spleen, bone marrow and blood of patients with ITP

Our cohort included 25 patients diagnosed with chronic or acute ITP, with a median age of 51 years (ranging 21 to 79 years), to investigate the anatomical distribution, affinity and secretion rate of anti-GPIIbIIIa IgG-SC. Clinical characteristics are presented in Table 1. Eight patients (8/18) achieved complete response after splenectomy, with a follow-up of 14-24 months, while 10 patients (10/18) had no significant increase in platelet counts after splenectomy. A bone marrow aspirate was also performed in 7/18 patients in addition to the programmed splenectomy. We also analyzed bone marrow and blood from 9 patients with ITP that were not splenectomized (Table 1), and twenty-one healthy donors (no immune disease) as controls for spleen, bone marrow or blood samples. For every sample, we analyzed an average of 50,000-100,000 droplets in total, representing an average of 10,000-27,000 single mononuclear cells containing 0,01-5% IgG-SC (Supplementary Figure S2). Compared to the detection of anti-GPIIbIIIa IgG-SC by ELISPOT, DropMap was far more sensitive (Supplementary Figure S3a).

**Table 1.**
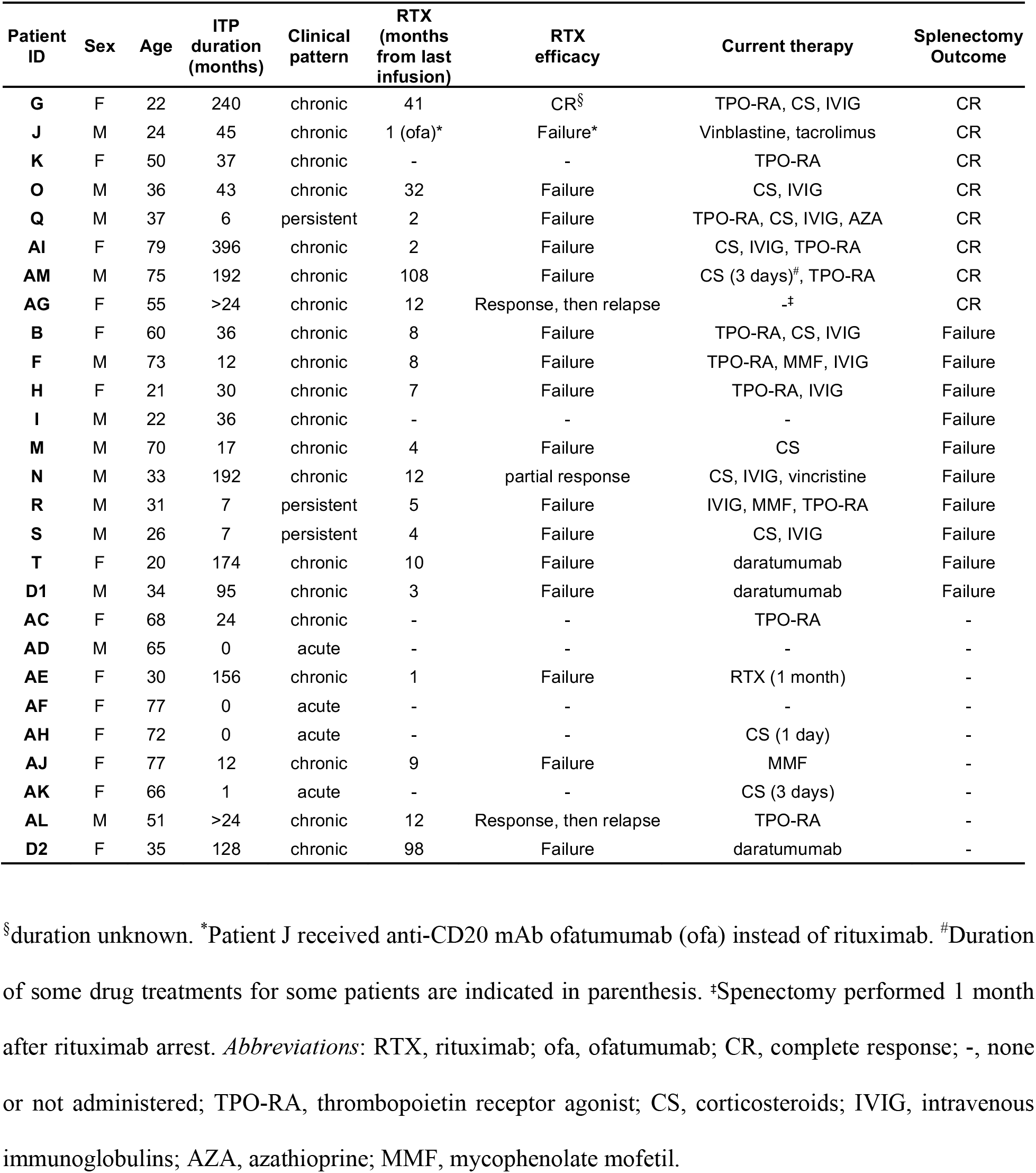
Characteristics of patients with ITP.

We first analyzed the global IgG-SC response per organ for all patients and healthy donors and found that among mononuclear cells 0.01% - 5% had detectable levels of IgG secretion, with a wide range of IgG secretion rates (1 - 713 IgG/s) (Figure 2a, Supplementary Figure S2). In healthy donors IgG secretion rates were not significantly different between the spleen, blood and bone marrow. However, in patients with ITP the IgG secretion rates in the bone marrow were significantly higher than in spleen and blood. Unexpectedly, splenic IgG-SC showed significantly (2.5-fold) lower secretion rates in patients with ITP compared to healthy donors, with median values of 46 IgG/s and 116 IgG/s, respectively. Similar findings were observed for IgG-SC from peripheral blood, with a 2-fold lower secretion rate in patients with ITP (median: 37 IgG/s) compared to healthy donors (74 IgG/s). However, IgG-SC in the BM had similar secretion rates between patients with ITP and healthy donors (median: 60 and 59 IgG/s, respectively) (Figure 2a). In all three compartments, the top ~20% highest IgG producers were responsible for ~50% of the total amount of secreted IgG (Supplementary Figure S4).

**Figure 2.**
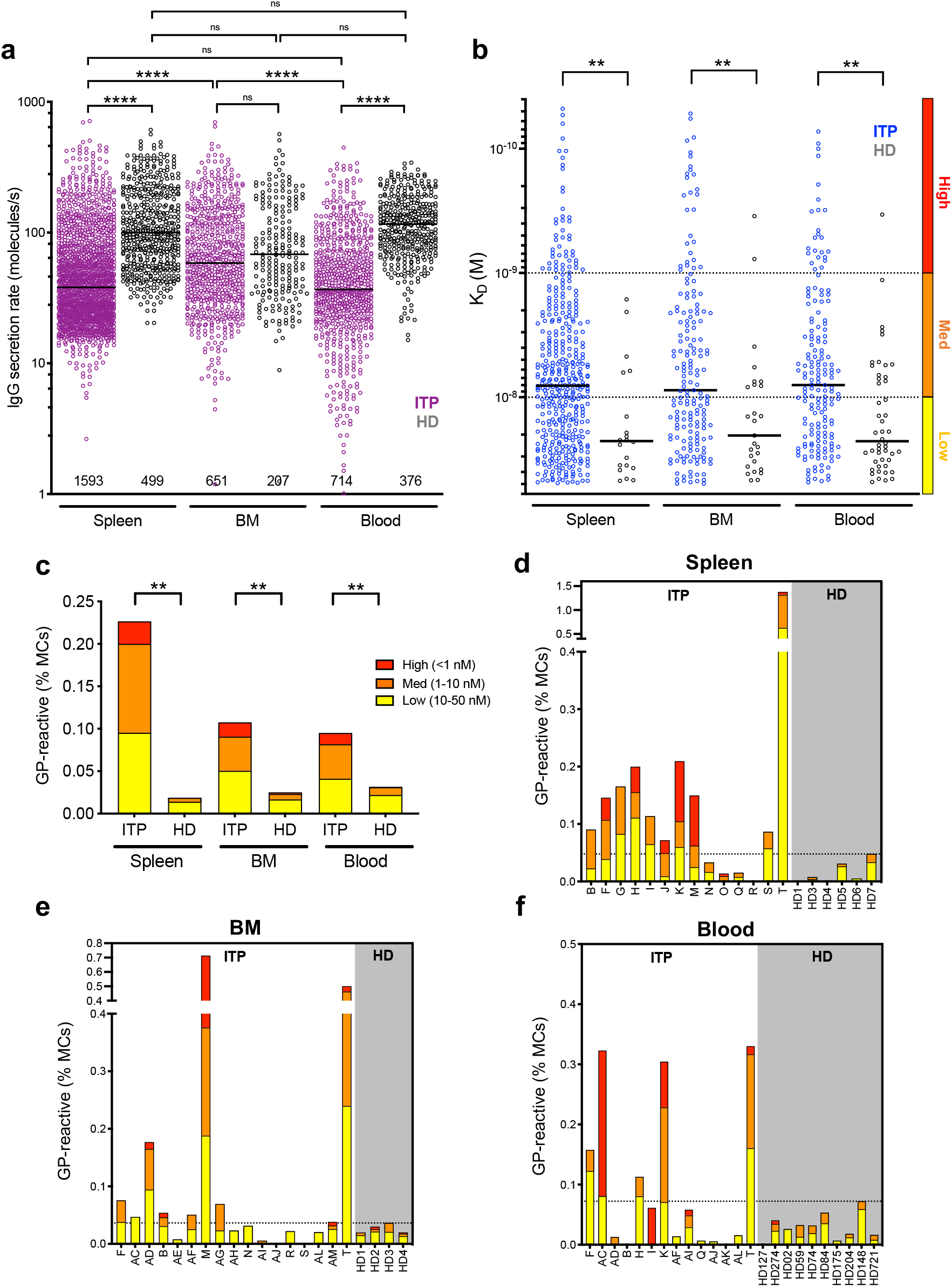
Autoreactive ASCs with high and low affinity are present in the spleen, blood and BM of patients with ITP. (**a**) IgG secretion rate and (**b**) affinity for GPIIbIIIa of single ASCs from pooled data of patients with ITP and healthy donors (HD) for spleen (ITP, n=14; HD, n=6), bone marrow (ITP, n=17; HD, n=4) and blood (ITP, n=14; HD, n=10). Single cell values and medians are plotted. Total number of IgG-SC cells analyzed per compartment are indicated in (a). In (b), affinities are classified into “high” (red), “medium” (orange) and “low” (yellow) affinity with dotted lines separating these categories. (**c**) Frequency of GPIIbIIIa-reactive IgG-SC among mononuclear cells (MCs) classified into “high”, “medium” and “low” affinity from patients (ITP) and healthy donors (HD) according to (b). (**d**-**f**) Frequency of GPIIbIIIa-reactive IgG-SC among mononuclear cells (MCs) classified into “high”, “medium” and “low” affinity represented for individual patients (ITP) and healthy donors (HD) for (**d**) spleen, (**e**) bone marrow (BM) and (**f**) blood. (d-f) The dotted line marks the highest frequency found in a healthy donor. NS (not significant); *P < 0.05; **P < 0.01; ***P < 0.001 using post-hoc contrast analysis after linear (a-b) or logistic (c) modelisation and multiple testing P value adjustment. See Supplemental Table 2 for further details.

Similar to the first patient we analyzed as a proof-of-concept (Figure 1c-e), we identified IgG-SC with affinity in the 5×10^−8^ M – 5×10^−11^ M range for GPIIbIIIa in pooled data from all three anatomical sites of patients with ITP, but also in significantly fewer numbers in pooled data from healthy donors in the 5×10^−8^ M – 5×10^−9^ M range (Figure 2b). Only patients with ITP had IgG-SC with very-high estimated affinity (K_D_ below 10^−10^ M), mathematically extrapolated from values outside the boundaries of the reference curve presented in Fig.1b. No correlation was found between affinity and secretion rate using the pooled data (Supplementary Figure S4). Only a fraction (8-13%) of IgG-SC showed crossbinding to GPIIbIIIa and antigens used for polyreactivity testing (keyhole limpet hemocyanin and insulin^24^) with low affinity (10-50nM), as expected from IgG-secreting plasma cells^25^, and increased affinity for GPIIbIIIa did not increase affinity for these other antigens (Supplementary Figure S3b-c). These results demonstrate that ~90% of the IgG-SC with reactivity for GPIIbIIIa analyzed herein are not polyreactive. The overall median affinity of anti-GPIIbIIIa IgG-SC from pooled patients with ITP was identical (~8 nM) between the three anatomical compartments (Figure 2b). The same result was found for anti-GPIIbIIIa IgG-SC of pooled healthy donors, with a significantly lower median affinity (~23 nM) (Figure 2b). The proportion of autoreactive IgG-SC among mononuclear cells was 10-fold, 5-fold and 3-fold higher in patients with ITP than healthy donors in the spleen, bone marrow and blood respectively (Figure 2c). Healthy donors harbored 65-75% low-affinity and relatively few high-affinity IgG-SC, whereas patients with ITP harbored 37-46% medium affinity and 11-16% high-affinity IgG-SC in the three anatomical compartments analyzed. Compared to healthy donors, patients with ITP secrete IgG at lower rate but with a higher proportion of medium- and high-affinity anti-GPIIbIIIa specificities in spleen, bone marrow and blood. These results suggest a role for high-affinity autoantibodies secreted from different anatomical locations in the pathogenesis of ITP.

We found varying proportions of autoreactive IgG-SC among mononuclear cells between patients with ITP, and within anatomical compartments of a given patient (Figure 2d-f). This finding may rely on the heterogeneity of therapeutic regimens and intrinsic cell heterogeneity of each anatomical compartment. Nevertheless, 71% (10/14) of spleen samples, but only 41% (7/17) of bone marrow and 36% (5/14) of blood samples harbored more autoreactive IgG-SC in patients with ITP compared to the highest value found in healthy control samples. 50% (7/14) of the patients with ITP harbored high-affinity anti-GPIIbIIIa IgG-SC in the spleen, compared to 29% (5/17) and 36% (5/14) in the bone marrow and blood, respectively. No such cell could be detected in spleens from healthy donors, and only once in blood and twice in bone marrow at very low numbers. Thus, most patients with ITP displayed a robust anti-GPIIbIIIa response in the spleen underlying the central role of this organ in ITP.

### Comparable autoantibody responses in paired organs

In order to compare the autoantibody response in different immune sites from the same patient with ITP, we obtained paired samples, either taken at the time of splenectomy (spleen + bone marrow and/or blood) or at the time of bone marrow collection (bone marrow + blood). On average 20-25% of IgG-SC were GPIIbIIIa-specific in the three compartments, with some patients presenting with a robust autoreactive response up to 75% of autoreactive IgG-SC among all IgG-SC (Figure 3a). Distribution of affinities towards GPIIbIIIa was similar within compartments of individual patients, as exemplified in Figure 3b (refer to Supplementary Figure S6 for all paired samples). For example, patient K displayed a robust anti-GPIIbIIIa response with >50% autoreactive IgG-SC distributed among high-, medium- and low-affinity IgG-SC in both spleen and blood; patient T displayed large numbers of IgG-SC, with ~25% autoreactive IgG-SC largely predominated by low- and medium-affinity IgG-SC in all three compartments; patient N displayed low numbers of IgG-SC, with ~5% autoreactive IgG-SC (Figure 3b). Generally, in the cohort of paired samples, patients harboring a large proportion of GPIIbIIIa-reactive IgG-SC among all IgG-SC in one compartment, also did so in the other one or two compartments (Figure 3c). For the 7 patients for whom splenectomy failed to induce clinical remission and a bone marrow sample was available (patients M, T, F, B, S, N, R), anti-GPIIbIIIa IgG-SC were present in the bone marrow the day of splenectomy, except for patient S. This suggests that the autoreactive bone marrow ASC population is responsible for the sustained disease observed after splenectomy in these patients.

**Figure 3.**
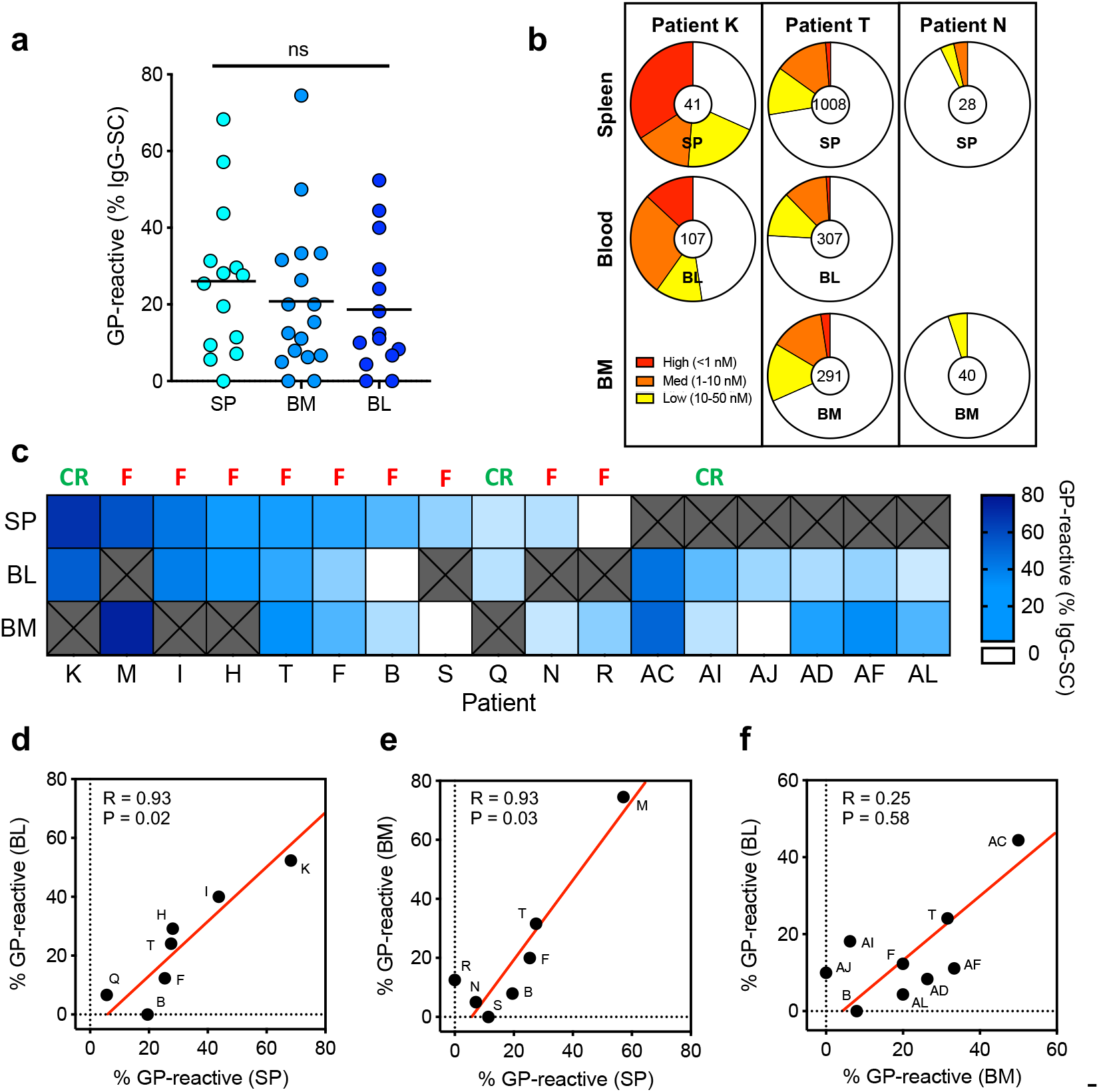
Paired organs demonstrate a comparable autoantibody response. (**a**) Frequency of GPIIbIIIa-reactive IgG-SC among total IgG-SC from individual patients with ITP. Each dot represents an individual. Values are pooled data from two replicates. NS (not significant); *P < 0.05; **P < 0.01; ***P < 0.001 using a Welch test and multiple testing P value adjustment. See Supplemental Table 2 for further details. (**b**) Distribution of IgG-SC into low (yellow), medium (orange) and high (red) affinity binders to GPIIbIIIa or non-binders (white), with total IgG-SC numbers indicated, for paired samples of patients K, T and N. (**c)** Heatmap of the frequency of GPIIbIIIa-reactive cells among IgG-SC per patient, ordered from most frequent to less frequent in spleen for patients with a spleen sample, and ordered from most frequent to less frequent in blood for patients without a spleen sample. White boxes indicate undetectable reactivity for GPIIbIIIa, and “x” indicate absence of samples. Result of splenectomy is indicated: complete remission (CR) or failure (F). (**d-f**) Pearson correlation analysis of data from (**c**); scatterplots compare the frequency of GPIIbIIIa-reactive cells among paired organs. R and adjusted P values are indicated. Red line indicates the Reduced Major Axis.

Proportions of GPIIbIIIa-reactive IgG-SC correlated well between spleen and blood (R>0.9; P=0.02) and spleen and bone marrow (R>0.9; P=0.03), but did not correlate between bone marrow and blood (R=0.25; P=0.58) (Figure 3d). Despite the heterogeneity of clinical situations and treatment history among patients in this cohort, the spleen appears to determine the extent of the autoreactive response, likely by providing the other compartments with autoreactive IgG-SC. Our results suggest that the autoreactive IgG response in patients with ITP disseminates through multiple organs, resulting in IgG-SC populations with similar ranges of anti-GPIIbIIIa affinity and proportion among IgG-SC.

### Kinetic follow-up of anti-CD38 therapy by daratumumab in patients with ITP

The autoreactive IgG-SC in patients with chronic ITP that are refractory to conventional therapies (rituximab and/or splenectomy) can be theoretically eliminated using the anti-CD38 monoclonal antibody daratumumab that was developed to target malignant plasma cells^26^. We therefore analyzed 3 patients who received daratumumab “off label” (compassionate use) for severe chronic refractory ITP. After three infusions, daratumumab led to an 89% reduction in circulating CD27^+^ P63^+^ plasmablasts and plasma cells^27^, identified by flow cytometry (Fig.4a; Supplementary Figure S5).

**Figure 4.**
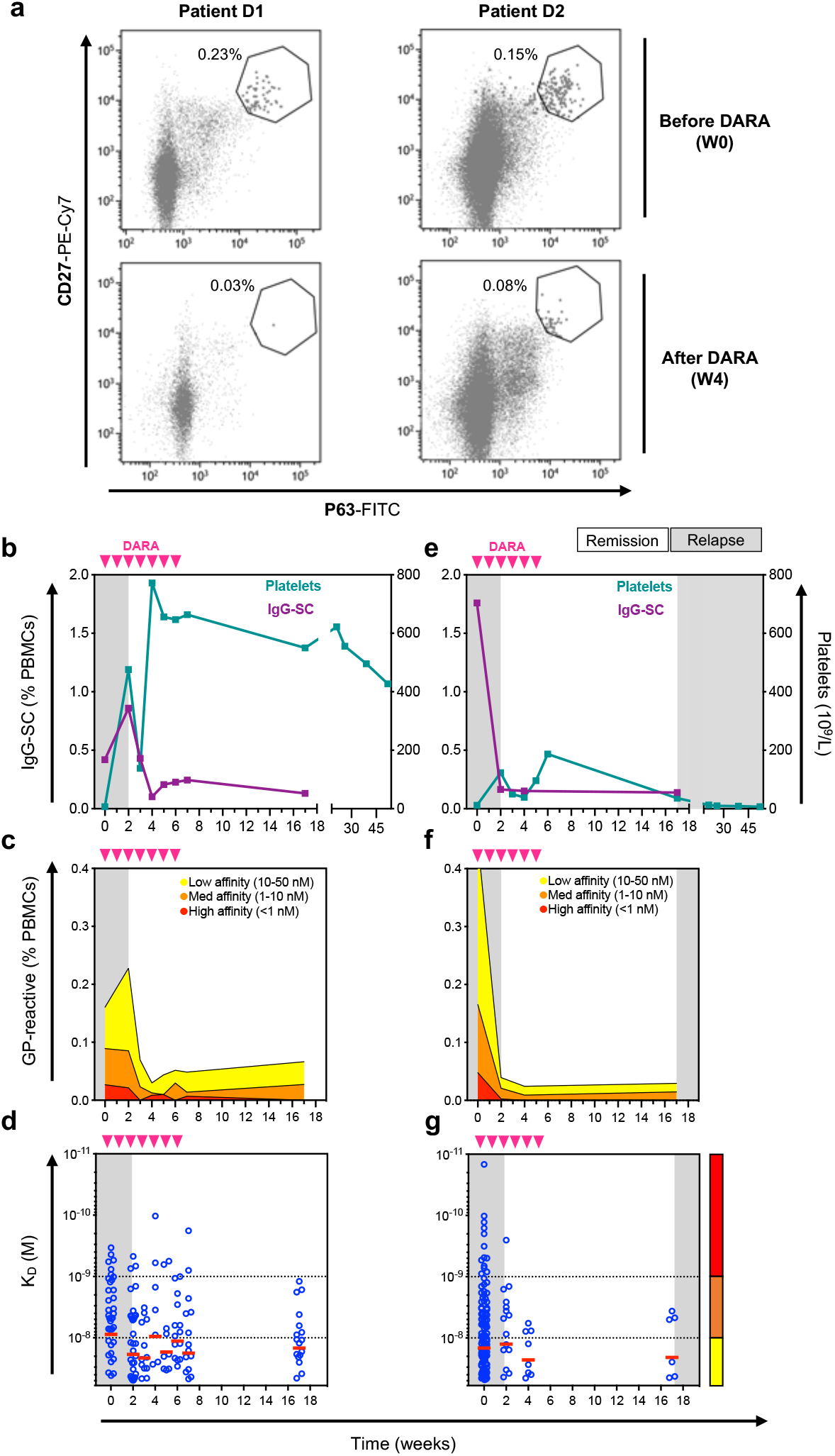
Anti-CD38 therapy deplete pathogenic ASC in patients with ITP. **(a)** Flow cytometric identification of circulating CD27^+^P63^+^ plasmablast/plasma cells from patients D1 and D2 before (top panel) and 4 weeks after (bottom panel) the start of daratumumab (DARA) treatment. (**b-g**) Kinetic follow-up of patient D1 (b-d) or patient D2 (e-g) blood samples for (**b,e**) frequency of IgG-SC among PBMCs and platelet levels, (**c,f**) frequency of GPIIbIIIa-reactive IgG-SC among PBMCs classified into “high” (red), “medium” (orange) and “low” (yellow) affinity, (**d,g**) affinity for GPIIbIIIa of single IgG-SC. Displayed on the background is time spent in clinical remission (white) or relapse (grey). Daratumumab infusions are indicated by pink arrows. Single-cell affinity values and medians are plotted in (d) and (g).

Patient #D1 (34 y/o male) had been splenectomized 9 years prior to daratumumab therapy, but continued to experience skin and/or mucosal bleedings because of low platelet counts (< 30 x 10^9^/L) despite several immunosuppressant drugs (including rituximab, mycophenolate mofetil and ciclosporin) and thrombopoietin receptor agonist (TPO-RA). He received 7 daratumumab infusions at 16 mg/kg per week without other treatment except oral Dexamethasone 20 mg before each infusion and achieved a complete response lasting now over 1 year (defined by platelet count >100 ×10^9^/L) (Figure 4b). After a short increase in total IgG-SC (Figure 4b) and anti-GPIIbIIIa IgG-SC (Figure 4c) numbers in blood, these proportions dropped 2-3-fold and remained low for at least 10 weeks after the last daratumumab infusion. Remarkably, whereas low-affinity IgG-SC remained at a third of their initial level, high-affinity IgG-SC became undetectable a few weeks after the end of the treatment (Figure 4c-d).

Patient #D2 (35 y/o female) had not been splenectomized and received her last rituximab infusion 8 years before receiving weekly daratumumab (without other treatment except oral Dexamethasone 20 mg before each infusion) for 6 weeks. This resulted in a transient complete response lasting 17 weeks before relapse. Total IgG-SC (Figure 4e) and anti-GPIIbIIIa IgG-SC numbers in blood nevertheless dropped ~10-fold following daratumumab infusions, again with the disappearance of high-affinity IgG-SC (Figure 4f-g). Overall, both patients responded similarly to daratumumab therapy in terms of depletion of IgG-SC, both total and GPIIbIIIa-specific, and elimination of high-affinity anti-GPIIbIIIa IgG-SC from circulation. These results emphasize that daratumumab therapy targets ASC in patients with ITP and suggest that the depletion of these cells, which include the autoreactive population, could be related to clinical improvement.

### Spleen-independent reappearance of high-affinity anti-GPIIbIIIa ASC after daratumumab

Patient T (20 y/o, female) had ITP requiring splenectomy 10 months after rituximab, then daratumumab therapy for refractory disease, allowing a sequential follow-up after these interventions. After splenectomy, IgG-SC and anti-GPIIbIIIa IgG-SC numbers in blood dropped 7- and 4-fold, respectively (Figure 5a,d), supporting the spleen as an important source of IgG-SC in circulation^21,28^. On the day of splenectomy, high numbers of IgG-SC were found among mononuclear cells in the blood (1.4%), bone marrow (1.6%) and spleen (5%) (Figure 5a-c) i.e., 7-fold higher than the ITP spleen with the highest numbers we had analyzed before (Supplementary Figure S2). Among IgG-SC, 27, 32 and 24 % were GPIIbIIIa-specific, including high-affinity IgG-SC, in the spleen, bone marrow and blood, respectively, demonstrating a relatively homogenous distribution of autoreactive IgG-SC across all three compartments (Figure 5d-i). Daratumumab treatment started 3-weeks after splenectomy and induced a transient increase in platelet counts (Figure 5a), associated with a decrease in total IgG-SC and anti-GPIIbIIIa IgG-SC numbers (Figure 5d) and leading to a disappearance of high-affinity IgG-SC (Figure 5g). Daratumumab was discontinued after 4 infusions for relapse and an increase of total IgG-SC, anti-GPIIbIIIa IgG-SC and high-affinity IgG-SC in the blood was observed after 5 weeks (Figure 5a,d,g). This reappearance of autoreactive, high-affinity IgG-SC in circulation occurred after splenectomy, suggesting autoreactive B cell reservoirs were present in other anatomical compartments or re-emergence of autoreactivity from naive B cells in secondary lymphoid organs (e.g., lymph nodes)^18^. Remarkably, whereas bone marrow IgG-SC (Figure 5c) and anti-GPIIbIIIa IgG-SC (Figure 5f) were reduced 2.5-3-fold compared to their content before splenectomy and daratumumab treatment, high-affinity IgG-SC could be readily detected in the bone marrow 5 weeks later with a similar distribution (Figure 5i). These bone-marrow resistant autoreactive high-affinity IgG-SC identified after multiple therapies could correspond to daratumumab-resistant ASC, and/or to newly immigrating ASC generated in another compartment that remains to be identified.

**Figure 5.**
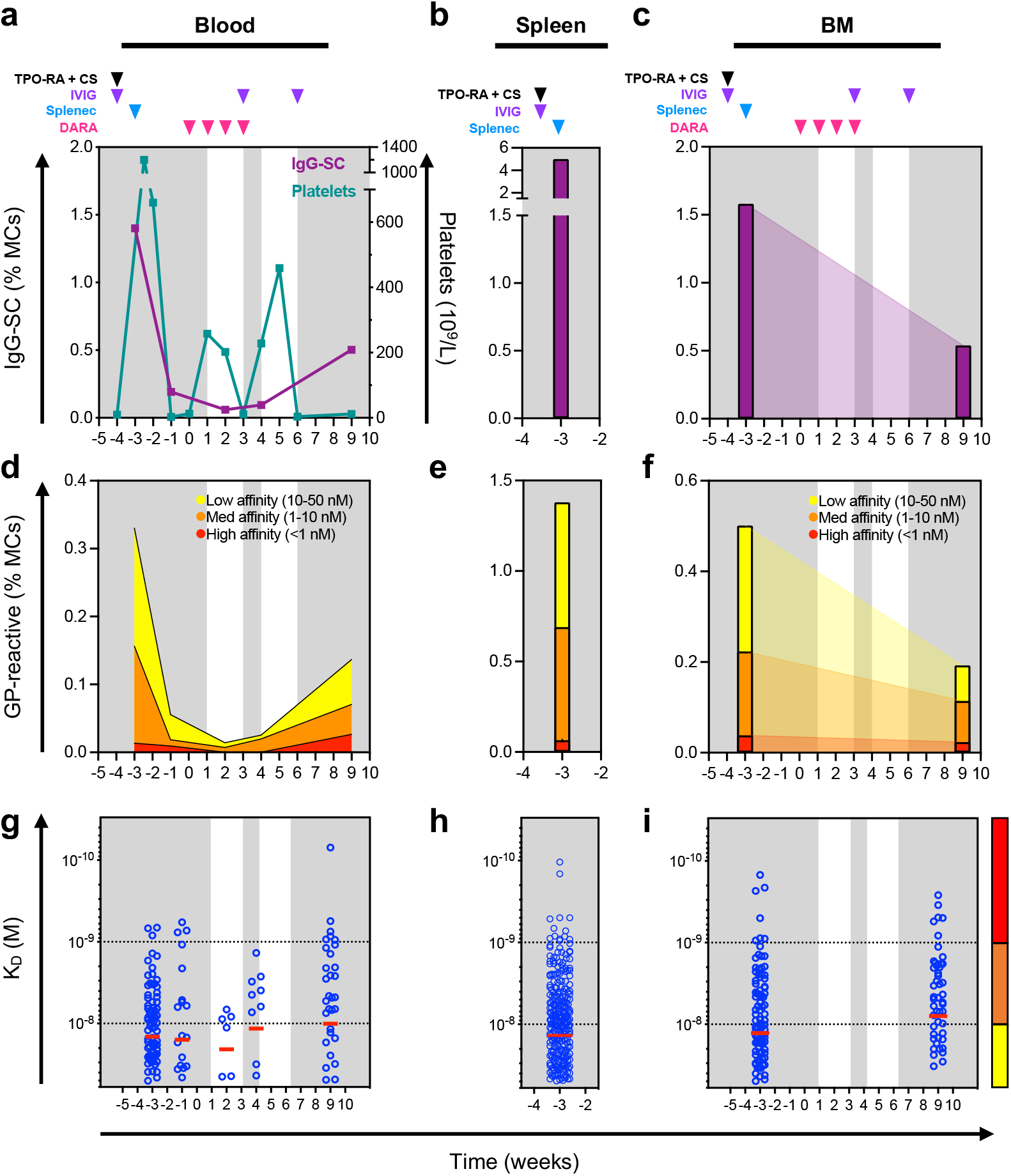
Failure of anti-CD38 treatment is associated with persistence of pathogenic ASC. Kinetic follow-up of patient T. Displayed on the background is time spent in clinical remission (white) or relapse (grey). Drug infusions are indicated by arrows: thrombopoietin receptor agonist and cyclosporin (TPO-RA + CS; black), intravenous immunoglobulin (IVIG; purple), daratumumab (DARA; pink). The time of splenectomy is indicated by a blue arrow. (**a-c**) Frequency over time of IgG-SC among mononuclear cells in (**a**) blood with platelet levels indicated, (**b**) spleen and (**c**) bone marrow. (**d-f**) Frequency over time of GPIIbIIIa-reactive IgG-SC among mononuclear cells in (**d**) blood, (**e**) spleen and (**f**) bone marrow classified into “high”, “medium” and “low” affinity. (**g-i**) Affinity over time for GPIIbIIIa of single IgG-SC in (**g**) blood, (**h**) spleen and (**i**) bone marrow. Single-cell affinity values (blue dots) and medians (red lines) are plotted.

## Discussion

This study represents the first characterization of autoantibody-secreting cells in humans in terms of their anatomical distribution, antibody secretion and affinity for their target autoantigen, including both patients with autoimmune disease and healthy donors. It demonstrates the dissemination of autoreactive IgG-secreting cells, among the spleen, blood and bone marrow from patients with ITP. Rates of IgG secretion per cell were very diverse, but globally similar between IgG-SC from the spleen and blood, but dissimilar from IgG-SC from the bone marrow. Whereas affinities for the platelet autoantigen GPIIbIIIa varied over 3 logs, the median affinity and range of affinities were strikingly similar between the three compartments, suggesting a common origin. Indeed, the autoreactive IgG-SC population phenotype (i.e., proportion, affinity range) of the spleen appears to imprint that of the blood and bone marrow, supporting a dominant role in ITP^9^. The persistence of autoreactive ASC in the bone marrow after failure of anti-CD20 B cell depletion, splenectomy, and anti-CD38 therapy represents a new layer of complexity and target for ITP therapy.

Chronic ITP is the most common hematologic indication for splenectomy^29^, making this major secondary lymphoid organ accessible for research. GPIIbIIIa-reactive memory B cells^13,18^ and IgG-SC (plasma cells)^9^ were identified in the spleen of patients with ITP, as well as increased GPIIbIIIa-reactive effector T cells^9^ and reduced Tregs^12^ numbers compared to healthy controls. The stage of B cell development at which tolerance against platelet antigens breaks down remains to be determined. The identification in small numbers of anti-GPIIbIIIa IgG-SC in all healthy donors in this work supports the hypothesis of a low-level autoreactivity against platelets in healthy individuals. This is in agreement with the low frequency (~13%) of self-reactive IgG-secreting plasma cells found in healthy donors after sorting of CD138^+^CD27^+^CD38^+^ cells and *in vitro* expression of their IgG^25^. As the abundance of anti-GPIIbIIIa IgG-SC is low, with poor affinity and IgG secretion rates, the amount and overall affinity of circulating platelet autoantibodies may be insufficient in these healthy donors to have a meaningful impact on platelet numbers, but may represent a biomarker for future evolution into ITP.

In ITP, tolerance breakdown against platelets lead to the generation of anti-GPIIbIIIa ASC clones that are a hallmark of germinal center autoreactive B cell generation^11,13,18^. We provided here a comprehensive view of affinity-matured autoreactive ASCs in different compartments that possess a broad range of affinities, including high-affinity binders. Autoantibodies with affinities for cytokines as high as 3×10^−14^ M have been reported to develop in APS1/APECED patients^30^. Whether antibody affinities for the extremely abundant antigen GPIIbIIIa could reach such values remains speculative and may certainly require an extreme defect in tolerance. Remarkably, anti-GPIIbIIIa subnanomolar affinities could be identified from IgG-SC in spleen, blood and bone marrow in similar distributions in some individuals. The magnitude of the response in the spleen correlated with that of the blood and bone marrow, with a strong decrease in circulating IgG-SC after splenectomy, highlighting the role of this organ in the generation of IgG-SC that are released to the bloodstream. Supportive of this notion, a correlation in the number of GPIIbIIIa-reactive IgG-secreting cells between spleen and blood from patients with ITP that responded to splenectomy was reported earlier using ELISpot^9^.

Only one case report identified previously anti-GPIIbIIIa in the bone marrow of one ITP patient using ELISpot^31^. Our study identified anti-GPIIbIIIa IgG-SC in the bone marrow in 40% of patients with ITP in similar or higher proportions among mononuclear cells than in the spleen. In a direct comparison, the sensitivity of DropMap was largely higher than ELISpot. This may be due to the inherent higher sensitivity of fluorescent bioassays compared to chromogenic assays, or to the necessity of IgG-SC to secrete over longer periods of time for ELISpot (hours) than for DropMap (45 min). The bone marrow is thought to be the main niche for survival of long-lived ASC, so the presence of autoreactive IgG-SC in this compartment could be correlated to long-term autoantibody production and, in turn, with the maintenance of chronic disease (reviewed in^32^). Our results constitute one of the first direct findings of autoreactive plasma cell populations in human autoimmunity^33^, supported also by similar findings in mouse autoimmunity models of vasculitis^34^, lupus^35–38^ and encephalomyelitis^39^. The establishment of long-lived ASC in the bone marrow could also have important implications for treatment of ITP and other B cell-dependent autoimmune disorders: first, this population could sustain autoantibody levels following splenectomy, which may be sufficient to affect clinical manifestations; second, long-lived ASC in the bone marrow have been observed to be resistant to immunosuppressive agents^36^ and to B-cell targeted therapies, such as anti-CD20 or anti-BAFF antibodies^32,39^, which may partly explain therapeutic failures.

In this context, targeting ASC with anti-CD38 appeared to be a very promising option. This work reports anti-CD38 daratumumab off-label use in three refractory chronic ITP patients, a treatment that has been proposed for autoimmune cytopenia following bone marrow transplantation^40,41^. Similar to our results, daratumumab therapy gave drastically different clinical outcomes, from complete remission to failure of therapy^41^. The bone marrow is probably responsible for immediate failures of splenectomy by harboring sufficient autoreactive ASC for effective platelet destruction. However, targeting autoreactive ASC may prove difficult as bone marrow cells have been reported in mouse models to be rather resistant to therapeutic antibody depletion^42,43^. Similarly, we showed that autoreactive ASC persisted in the bone marrow of a patient with no response to daratumumab, providing a possible explanation for treatment resistance. Another patient achieved complete remission after daratumumab but relapsed several weeks after, suggesting that a lymphoid organ was perhaps able to generate new autoreactive ASC. The lymphoid organ responsible may have been lymph nodes in the splenectomized patient T (Figure 5) and the spleen (and perhaps also lymph nodes) in patient D2 who received daratumumab but had not been splenectomized (Figure 4e-g). Understanding the reasons for such variability in clinical responses to B cell depletion therapy will require more investigation, particularly as we describe herein a relatively homogenous dissemination of IgG-SC among spleen, blood and bone marrow of patients with ITP.

Our work also has several limitations, mostly inherent to human studies. Patients had varying clinical histories and received treatments that probably impacted the ASC pool. However, long-term corticosteroids and/or immunosuppressant drugs are not commonly given for ITP in France, and we took advantage of the various therapeutic sequences including splenectomy and daratumumab to study ASC dissemination in human. Our assay focused on IgG and GPIIbIIIa as previous studies showed that anti-GPIIbIIIa IgG were predominant autoantibodies in ITP^4,5^. We cannot, however, exclude that some patients had a response of another isotype or directed against other, less common, platelet antigens (e.g. GPIb-IX and GPIa-IIa)^44^. ASC are also rare cells among mononuclear cells, and in some patients described here only few anti-GPIIbIIIa reactive cells could be identified that limit broad extrapolations on autoreactive ASC. Our novel method of analysis proved nevertheless more sensitive than conventional assays (i.e., ELISpot), which allowed us to investigate secretion rate and affinity for GPIIbIIIa for >3,300 freshly isolated *ex vivo* IgG-SC.

This work extends^45–48^ our knowledge of autoreactive IgG-SC in human ITP. We demonstrate the homogenous dissemination of platelet autoreactive ASC into multiple anatomical compartments that could explain splenectomy failure in some patients. A wide range of affinities for the major ITP autoantigen GPIIbIIIa were identified, with very high affinities reminiscent of affinities found in other systemic autoimmune diseases. Anti-CD38 daratumumab therapy allowed us to demonstrate the crucial contribution of ASC in ITP, and to point towards another compartment than the spleen that may serve in some patients as a source to reconstitute autoreactive ASC pools in circulation and in the bone marrow.

## Acknowledgements

The authors acknowledge Ms Laetitia Languille for the coordination of patient recruitment at Hôpital Henri Mondor, Créteil, France. P.C-H. was supported partly by a stipend from the Pasteur - Paris University (PPU) International PhD program, and by a fellowship from the French *Fondation pour la Recherche Médicale* (FRM). M.B. is the recipient of a CIFRE PhD fellowship. C.C. was supported by CONCYTEC, Peru. P.B. acknowledges funding from the French National Research Agency grant ANR-14-CE16-0011 project DROPmAbs, by the Institut Carnot Pasteur Microbes et Santé, the Institut Pasteur and the Institut National de la Santé et de la Recherche Médicale (INSERM). E.K. acknowledges funding from the Branco Weiss Fellowship (Society in Science) and the European Research Council (ERC) (grant agreement #80336). J.B. acknowledges funding from the French government through BPIFrance under the frame “Programme d’Investissements d’Avenir” (CELLIGO Project), the “Institut Pierre-Gilles de Gennes” through the laboratoire d’excellence, “Investissements d’avenir” programs ANR-10-IDEX-0001-02 PSL, ANR-10- EQPX-34 and ANR-10-LABX-3.

## Author Contributions

Experimental design, P.C-H, K.E, J.B, M.M and P.B; Investigation, P.C-H, E.C, M.B, A.S, G.C, I.A, A.V, A.P, B.I, O.R-L, C.C, G.M and D.S; Formal analysis, P.C-H, M.B, P.E, G.M, E.K, J.B, M.M and P.B; Writing (original draft), P.C-H and P.B; Writing (review and editing), all authors.

## Competing Interests statement

K.E. and J.B. have filed patents on the DropMap technology, and the inventors may receive payments related to exploitation of these under their employer’s rewards-to-inventors scheme. M.M. received research funds from GSK, outside of the submitted work and personal fees from LFB and Amgen, outside of the submitted work. P.B. received consulting fees from Regeneron Pharmaceuticals, outside of the submitted work. All other authors declare no competing interests.

---

**Figure S1.**
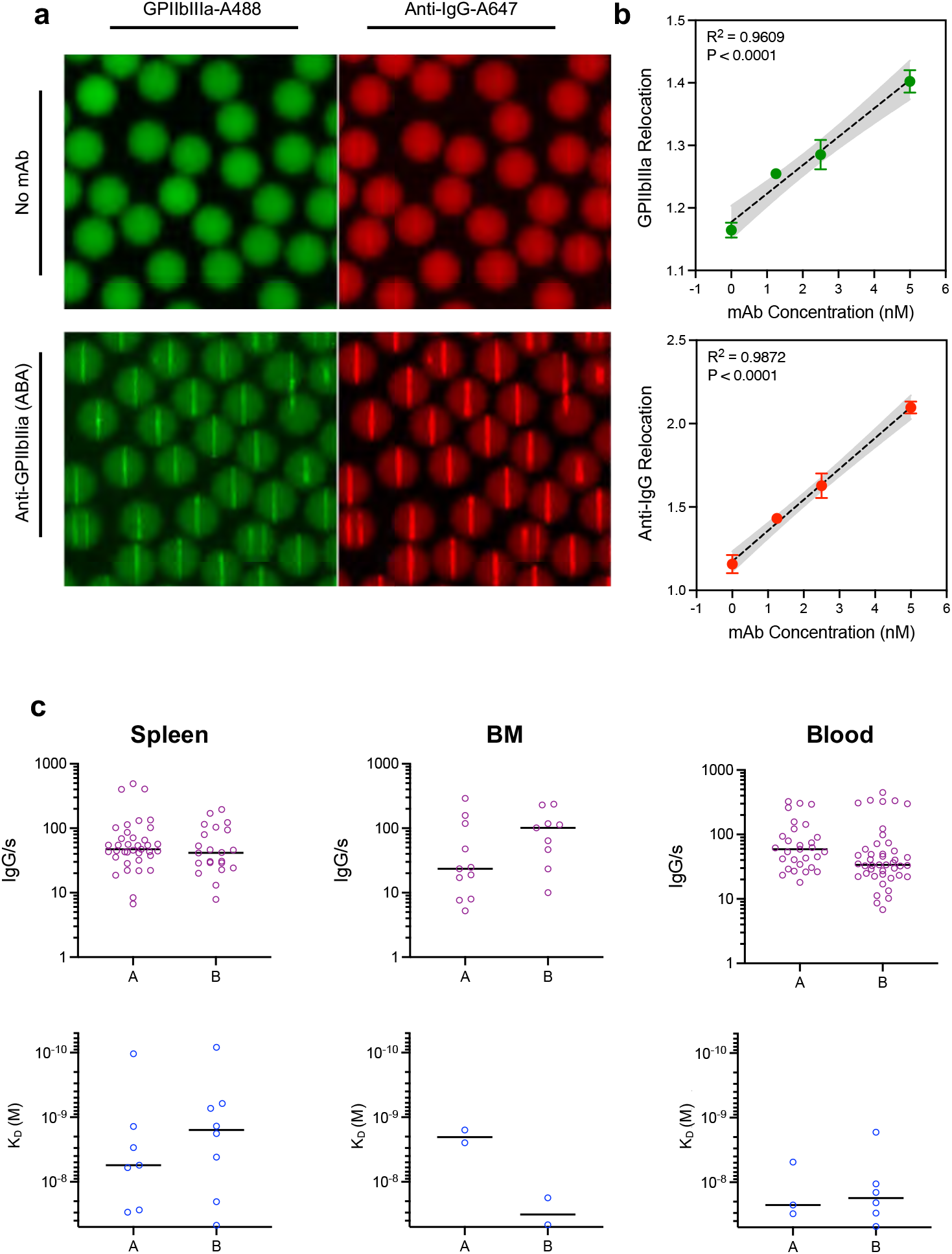
Bioassay calibration and reproducibility. (**a**) Representative images of droplet arrays containing a monoclonal antibody specific for GPIIbIIIa. (**b**) Quantification of fluorescent relocation in response to antibody concentration for GPIIbIIIa-Alexa488 (green) and anti-IgG F(ab’)2-Alexa647 (red). Fluorescent relocation is obtained by dividing the fluorescent signal from the beadline by the average fluorescent background signal of the drop. Relocation values for a range of monoclonal antibody concentrations were plotted and linear regression was performed. R-squared and P values are indicated. (**c**) Replicates for single-cell measurements. Data for IgG-secretion (top) and affinity for GPIIbIIIa (bottom) is shown for two replicates (A and B). All samples described in this work were analyzed in duplicate or triplicate.

**Figure S2.**
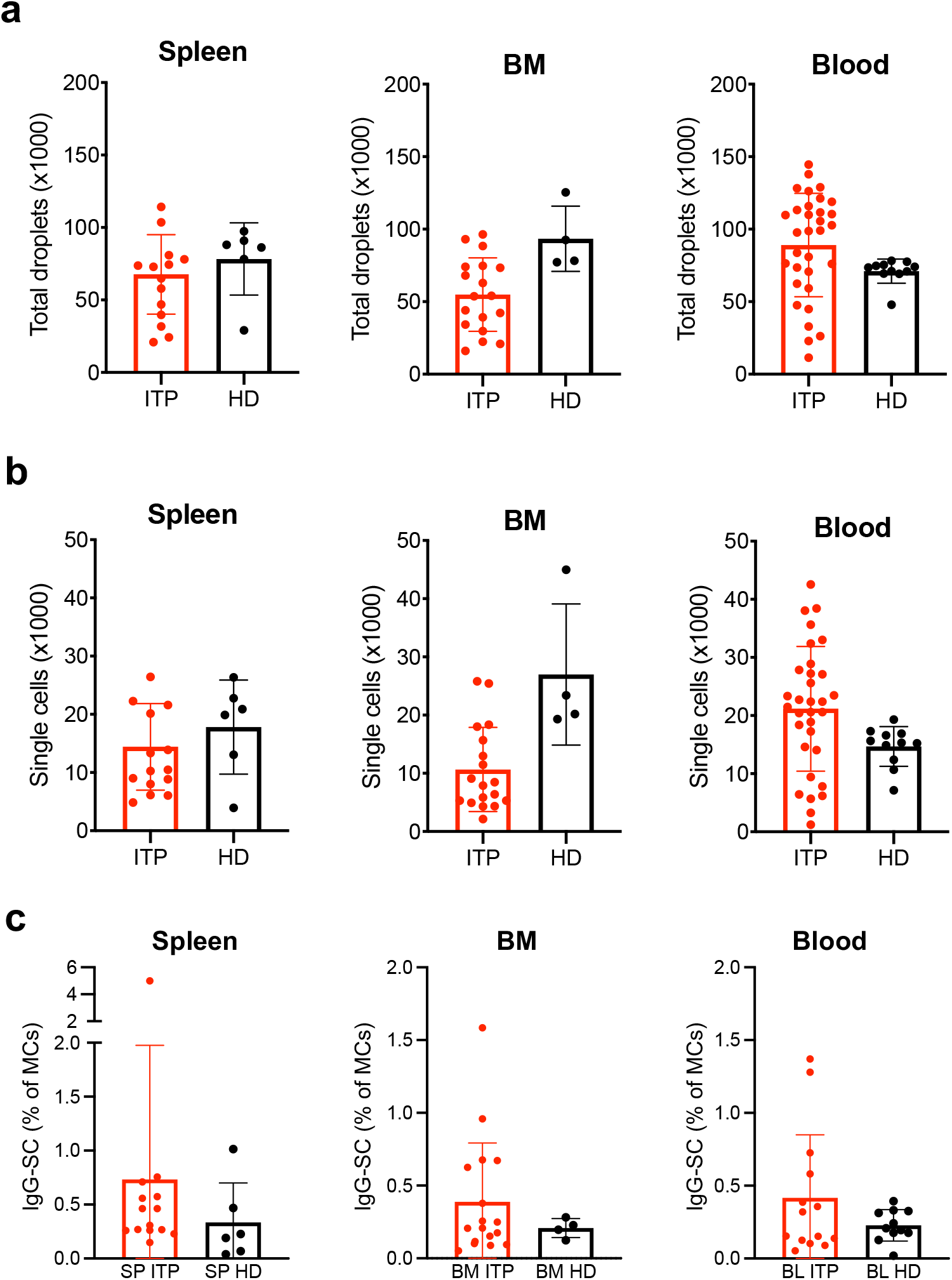
DropMap system throughput and IgG production by single cells. (**a**) Total number of droplets analyzed, (**b**) total number of single cells included in the analysis, and (**c**) number of IgG-SC found in each sample are represented for every sample described in this work. Every data point corresponds to a single sample after pooling of 2-3 replicate acquisitions. Data is shown as mean and SD.

**Figure S3.**
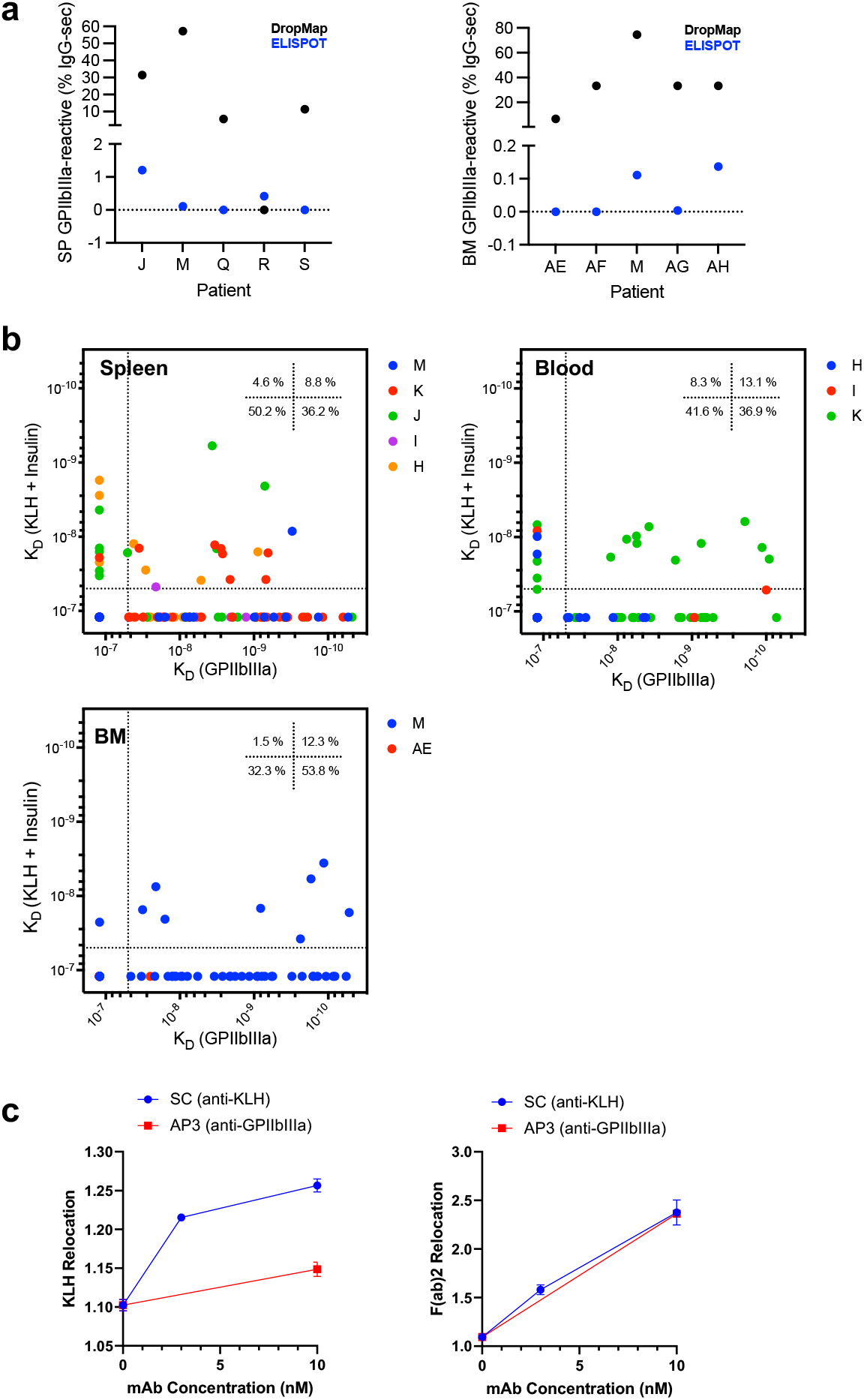
Sensitivity of the bioassay and polyreactivity measurements. (**a**) Comparison of percentage of GPIIbIIIa-reactive cells among IgG-SC identified by DropMap (black dots) or by ELISPOT (blue dots) in spleen (SP; left) and bone marrow (BM; right). (**b**) Affinity values of single IgG-SC against GPIIbIIIa and (KLH+insulin) from a multiplexed bioassay containing fluorescently labeled GPIIbIIIa-Alexa488, KLH-Alexa405 and Insulin-Alexa405. KLH and human insulin are labeled with the same fluorochrome to obtain a single affinity measurement [K_D_ (KLH+insulin)] for both these irrelevant antigens as a measure of polyreactivity. Distribution per quadrant is indicated with color codes per patient. (**c**) Positive and negative control for KLH binding using an anti-KLH IgG mAb (clone SC) and an anti-GPIIbIIIa IgG mAb (clone AP3) using the polyreactivity bioassay described in (b). Relocation of KLH (left) and anti-IgG are shown at different concentrations of the mAbs.

**Figure S4.**
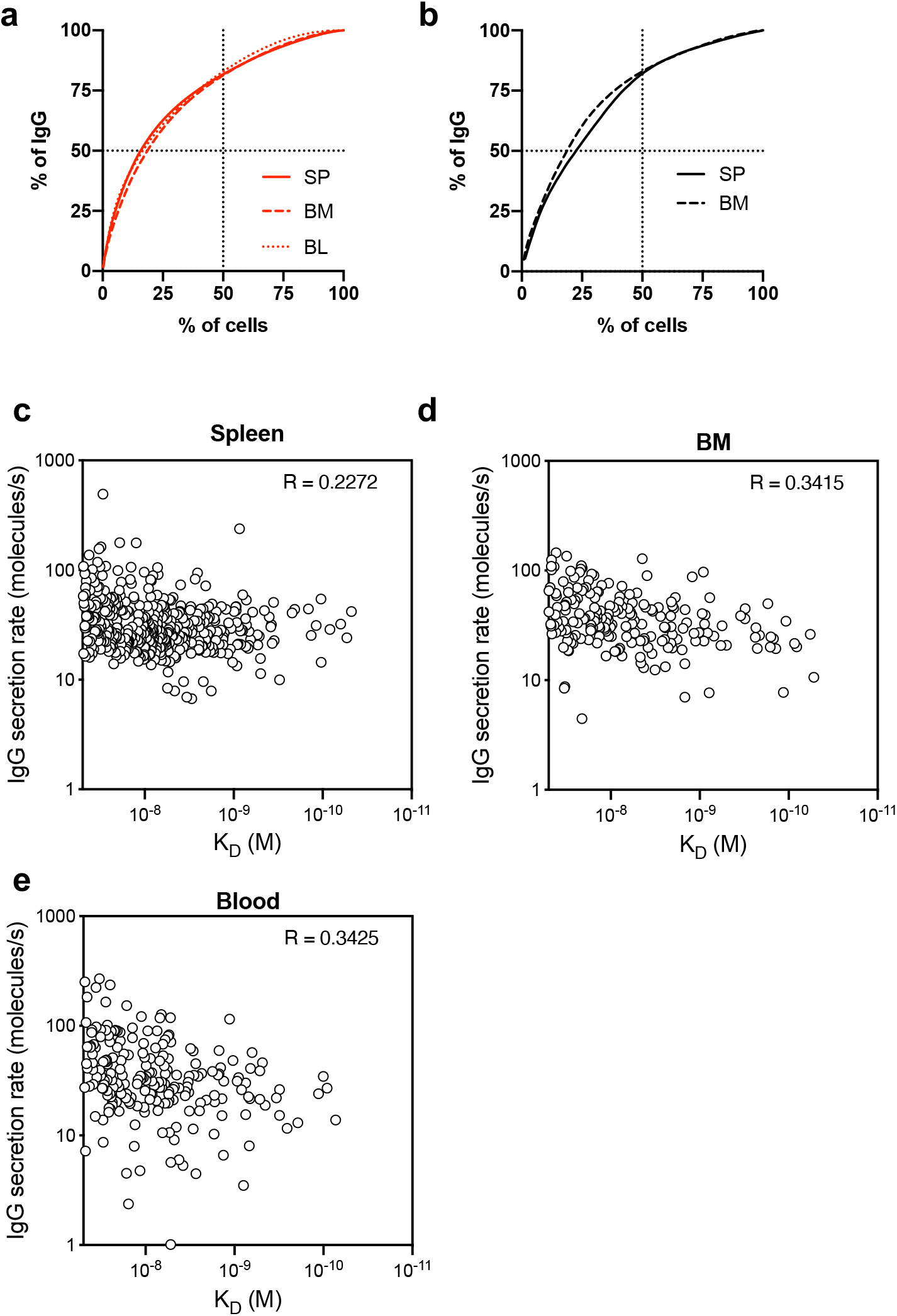
Cumulative IgG secretion and absence of correlation between IgG secretion and affinity for GPIIbIIIa. (**a-b**) Percentage of the total mass of secreted IgG relative to the percentage of IgG-SC ordered from highest to lowest IgG producer. Data for all the IgG-SC from all (a) patients with ITP and (b) healthy donors is pooled by organ. (**c-e**) Absence of correlation between IgG secretion rate and K_D_ for GPIIbIIIa. Pooled data from all anti-GPIIbIIIa IgG-SC identified from all (c) spleen, (d) bone marrow and (e) blood from all patients with ITP is shown. Correlation analysis performed using Pearson’s coefficient, R values are indicated.

**Figure S5.**
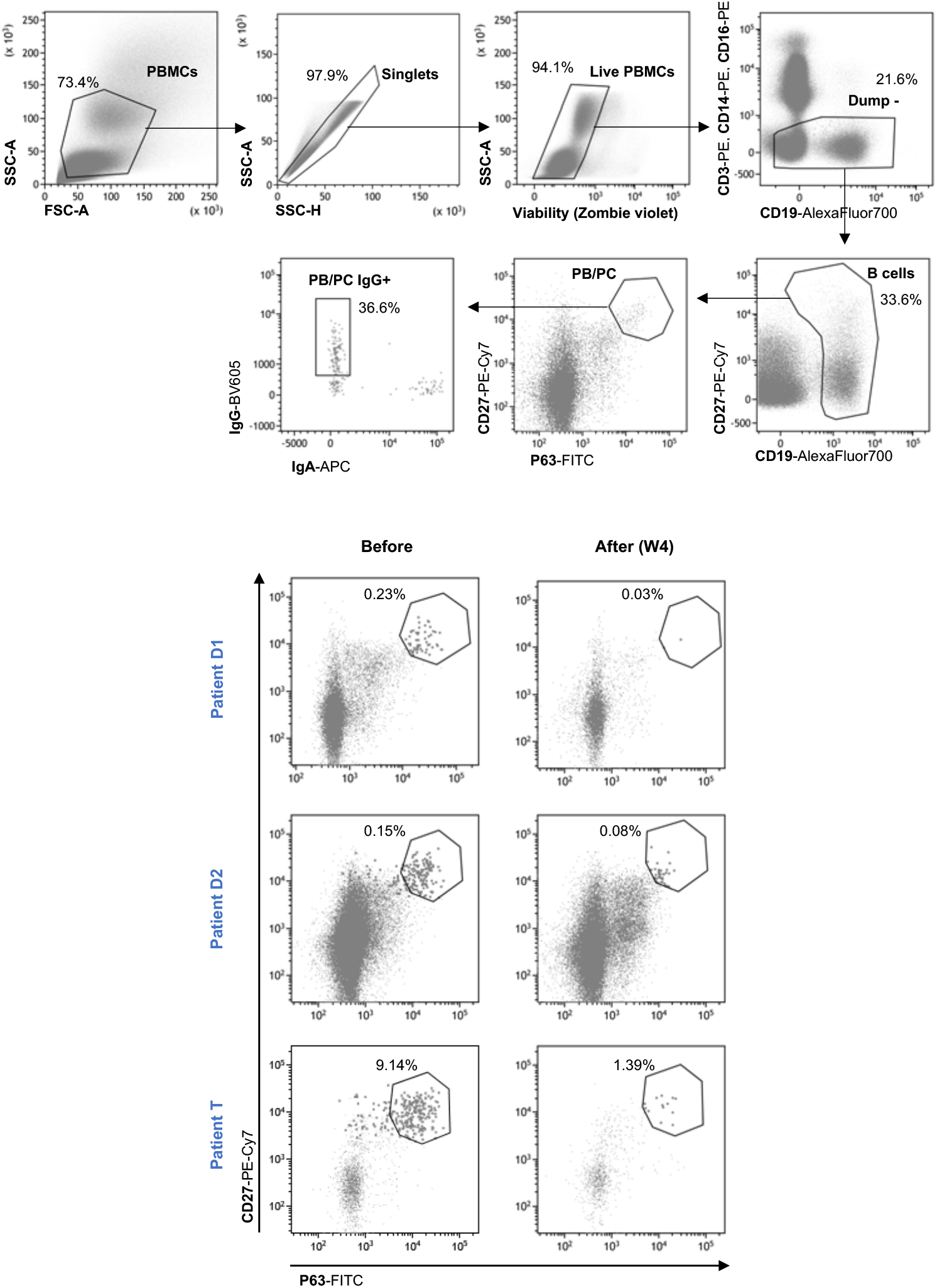
Identification of circulating plasma cells following daratumumab therapy. (Top) Gating strategy is shown for IgG^+^ plasma cell for a representative PBMC sample after Ficoll gradient and B cell enrichment by negative selection of CD3^+^ cells. Plasma cells are identified as live CD3^−^CD14^−^CD16^−^ CD19^+/−^CD27^+^p63^+^ cells and IgG^+^ cells. Percentages of cells within each gate are indicated. All antibodies used for this strategy are included in Supplementary Table 1. (Bottom) Follow-up of circulating IgG^+^ plasma cells (within the gate) before daratumumab therapy and at week 4 after the first infusion of daratumumab in patients D1, D2 (same data and dot plots as in Fig.4a) and T. Percentages of cells within each gate are indicated.

**Figure S6.**
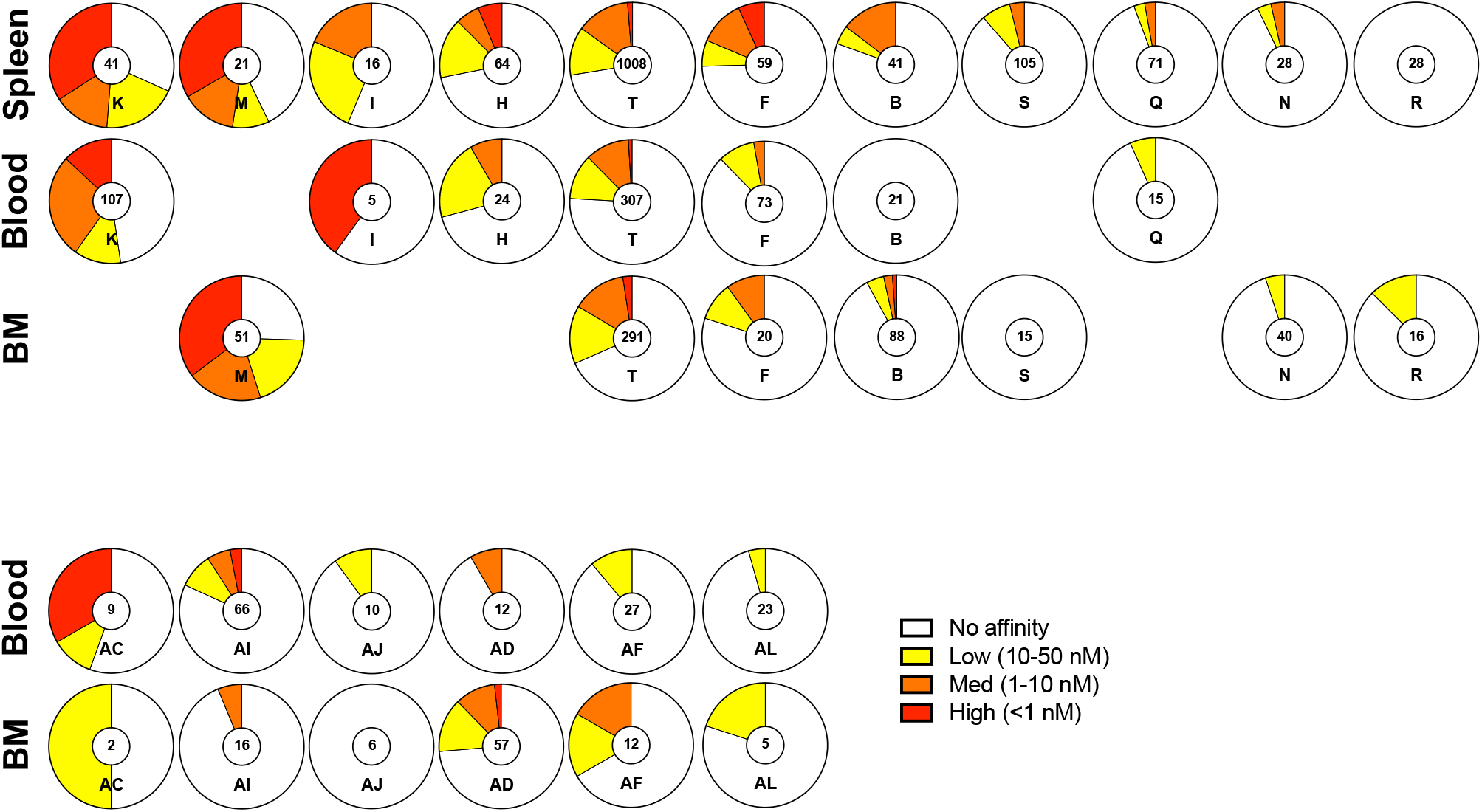
Affinity distribution among paired organs. Distribution of IgG-SC into low (yellow), medium (orange) and high (red) affinity binders to GPIIbIIIa or non-binders (white), with total IgG-SC numbers indicated, for all patients with ITP with paired samples in this study. Samples are represented ordered by the proportion of GPIIbIIIa-reactive cells present in the spleen (top) or blood (bottom).

**Table S1.** Reagents and software used for flow cytometry.

| Reagent | Source | Identifier | Fluorochrome |
| --- | --- | --- | --- |
| **Antibody for surface staining** |  |  |  |
| CD3 | BD Bioscience | UCHT1 ; 555333 | PE |
| CD14 | BD Bioscience | M5E2 ; 564054 | PE |
| CD16 | BD Bioscience | 3G8 ; 560995 | PE |
| CD38 | BD Bioscience | HIT2 ; 551400 | PerCP Cy5.5 |
| CD27 | BD Bioscience | M-T271 ; 560609 | PE-Cy7 |
| CD19 | BD Bioscience | HIB19 ; 557921 | AF700 |
| IgD | BD Bioscience | IA6-2 ; 562540 | PE-CF594 |
| **Antibody for intracellular staining** |  |  |  |
| VS38c | Dako | VS38c ; F7149 | FITC |
| IgG | BD Bioscience | G18-145 ; 563246 | BV605 |
| IgA | Miltinyi Biotec | IS11-8E10 ; 130-113-472 | APC |
| IgM | BD Bioscience | G20-127 ; 563113 | BV510 |
| **Chemical** |  |  |  |
| eBioscience™ Foxp3 / Transcription Factor Staining Buffer Set | ThermoFisher | 00-5523-00 |  |
| Zombie Violet™ Fixable Viability Kit | Biolegend | 423114 |  |
| Softwares |  |  |  |
| Kaluza v2.1 | Beckman Coulter |  |  |
| GraphPad Prism v8 | GraphPad |  |  |

**Table S2.** Statistical analyses.

| Figure | Comparison | Alternative hypothesis | Sample size | Transformation | Test | p-value | Adjusted p-value |
| --- | --- | --- | --- | --- | --- | --- | --- |
| 1C | SP and BM | mean difference | n = 59, n = 20 | log | Welch test | 0.76 | 0.97 |
|  | SP and BL | mean difference | n = 59, n = 73 | log | Welch test | 0.64 | 0.97 |
|  | BM and BL | mean difference | n = 20, n = 73 | log | Welch test | 0.97 | 0.97 |
| 1D | SP and BM | mean difference | n = 15, n = 4 | log | Welch test | 0.33 | 0.50 |
|  | SP and BL | mean difference | n = 15, n = 9 | log | Welch test | 0.01 | 0.04 |
|  | BM and BL | mean difference | n = 4, n = 9 | log | Welch test | 0.50 | 0.50 |
| 2A | ITP and HD in SP | effect difference | n = 1592, n = 399 \| df = 44.9 | log | Contrast test in linear model | 6e-6 | 1e-5 |
|  | ITP and HD in BM | effect difference | n = 706, n = 206 \| df = 38.8 | log | Contrast test in linear model | 0.63 | 0.70 |
|  | ITP and HD in BL | effect difference | n = 712, n = 376 \| df = 43.7 | log | Contrast test in linear model | 1e-7 | 4e-7 |
|  | SP and BM in ITP | effect difference | n = 1592, n = 706 \| df = 3787.7 | log | Contrast test in linear model | 5e-17 | 4e-16 |
|  | SP and BL in ITP | effect difference | n = 1592, n = 712 \| df = 3882.1 | log | Contrast test in linear model | 0.27 | 0.34 |
|  | BM and BL in ITP | effect difference | n = 706, n = 712 \| df = 3785.0 | log | Contrast test in linear model | 6e-16 | 3e-15 |
|  | SP and BM in HD | effect difference | n = 399, n = 206 \| df = 40.2 | log | Contrast test in linear model | 0.06 | 0.10 |
|  | SP and BL in HD | effect difference | n = 399, n = 376 \| df = 43.3 | log | Contrast test in linear model | 0.79 | 0.79 |
|  | BM and BL in HD | effect difference | n = 206, n = 376 \| df = 38.1 | log | Contrast test in linear model | 0.07 | 0.10 |
| 2B | ITP and HD in SP | effect difference | n = 1592, n = 399 \| df = 40.4 | log | Contrast test in linear model | 0.005 | 0.005 |
|  | ITP and HD in BM | effect difference | n = 706, n = 206 \| df = 34.2 | log | Contrast test in linear model | 0.005 | 0.005 |
|  | ITP and HD in BL | effect difference | n = 712, n = 376 \| df = 38.6 | log | Contrast test in linear model | 0.005 | 0.005 |
| 2C | ITP and HD in SP | effect difference | n = 1592, n = 399 \| df = Inf | log | Contrast test in linear model | 0.006 | 0.006 |
|  | ITP and HD in BM | effect difference | n = 706, n = 206 \| df = Inf | log | Contrast test in linear model | 0.006 | 0.006 |
|  | ITP and HD in BL | effect difference | n = 712, n = 376 \| df = Inf | log | Contrast test in linear model | 0.006 | 0.006 |
| 3A | SP and BM | mean difference | n = 14, n = 17 | none (0 values) | Welch test | 0.4694 | 0.7041 |
|  | SP and BL | mean difference | n = 14, n = 14 | none (0 values) | Welch test | 0.2987 | 0.7041 |
|  | BM and BL | mean difference | n = 17, n = 14 | none (0 values) | Welch test | 0.7416 | 0.7416 |
| 3D-F | SP and BL | Correlation different from zero | n = 6 | log (0 values removed) | Pearson correlation test | 0.007 | 0.02 |
|  | SP and BM | Correlation different from zero | n = 5 | log (0 values removed) | Pearson correlation test | 0.02 | 0.03 |
|  | BM and BL | Correlation different from zero | n = 7 | log (0 values removed) | Pearson correlation test | 0.58 | 0.58 |
| S1-B | mAb and Relocation (GPIIb/IIIa) | Slope different from zero | n = 4 | none | Simple linear regression | 0.000019 | - |
|  | mAb and Relocation (IgG) | Slope different from zero | n = 4 | none | Simple linear regression | 6.6E-07 | - |

<sup>a</sup> n, sample size for each class tested, respectively; df, degree of freedom; adjusted p values from the same panel, according to Benjamini & Hochberg

## METHODS

### ITP patients and controls

All patients included in this study were adults (>18 y/o) and were diagnosed with ITP according to the international guidelines^1^. Patients with underlying immunodeficiencies, hepatitis C virus or human immunodeficiency virus infection, lymphoproliferative disorders, and defined systemic lupus erythematosus (≥4 American Rheumatism Association criteria) were excluded. Patients had acute (<3 months), persistent (3-12 months) or chronic (>12 months) ITP. Complete response to splenectomy was defined by a platelet count over 100_×_10^9^/liter and the absence of bleeding, partial response was defined by a platelet count over 30_×_10^9^/liter and below 100_×_10^9^/liter and at least doubling values from baseline, and failure by a platelet count under 30_×_10^9^/liter or use of salvage therapy after one month. Relapsing patients were defined as those that initially had a complete response, but then had a drop in the platelet count, below 30_×_10^9^/liter, as well as a medical intervention by the treating physician. Healthy controls (HDs) were healthy individuals with no autoimmune disease or lymphoma. Control spleen samples were obtained from organ donors that died from stroke or head trauma. Control bone marrow aspirate samples were taken from healthy organ transplantation donors and were obtained from the Pitié-Salpêtrière Hospital (AP-HP). Control blood samples were obtained from the French Blood Establishment (EFS).

This study was conducted in compliance with the Declaration of Helsinki principles and was approved by the Agence de la Biomédecine and the Institutional Review Boards Comité de Protection des Personnes (CPP) Ile-de-France IX (patients with ITP) and Ile-de-France II (HD spleen). All patients with ITP provided written informed consent before the collection of samples.

### Sample processing

Splenic tissue was obtained from splenectomy and maintained at 4°C for transportation to the laboratory. Splenic tissue fragments were homogenized using a dissociator (gentleMACS, Miltenyi), and cell suspensions were subsequently filtered and diluted in RPMI-1640 (Invitrogen). Ficoll density gradient (Ficoll Paque Plus, GE Healthcare) centrifugation at 1300 rpm for 20 min with no breaks was used to obtain Mononuclear cells (MCs). Bone marrow cells were obtained from bone marrow aspiration and immediately placed in sodium heparin tubes (BD) for transportation to the lab at room temperature. Bone marrow cells were then dissociated by flushing thoroughly through a syringe needle and diluted in RPMI-1640 (Invitrogen), before processing by Ficoll density gradient to obtain MCs. Blood was obtained in sodium heparin tubes and transported at room temperature. After dilution in RPMI, blood was processed by Ficoll density gradient. After isolation, MCs from all organs were resuspended in RPMI supplemented with 10% HyClone FBS (Thermo Scientific) and 1% Pen/Strep (Thermo Fisher). From this point on, cell suspensions were maintained on ice. All experiments were performed within 12 hours from sample collection.

### Aqueous phase I: preparation of cells for droplet compartmentalization

Cell suspensions were centrifuged (300g, 5min) and resuspended twice using MACS buffer consisting of PBS pH 7.2, 0.2% bovine serum albumin and 2 mM EDTA. After each resuspension, cells were filtered through a 40-μm cell strainer to eliminate aggregates. Cells were then spun (300g, 5 min) and resuspended in DropMap buffer, consisting of RPMI (without phenol red) supplemented with 0.1% Pluronic F68, 25mM HEPES pH 7.4, 5% KnockOut™ serum replacement (All Thermo Fisher), 0.5% human serum albumin (Sigma Aldrich). Cell number in the suspension was adjusted to achieve a λ (mean number of cells per droplet) of ~0.3. For calibration curves, monoclonal IgG antibodies were diluted in DropMap buffer.

### Aqueous phase II: preparation of beads and bioassay reagents

Paramagnetic nanoparticles (Strep Plus 300nM, Ademtech) were washed with Dulbecco’s phosphate-buffered saline with calcium and magnesium (DPBS++, ThermoFisher). The nanoparticles were resuspended in DPBS++ containing 1μM biotin-labeled anti-human kappa light chain (Igκ) (CaptureSelect; Thermo Fisher), and incubated 20min at room temperature. After another wash with DPBS++, nanoparticles were resuspended in 5% pluronic F127 (ThermoFisher) and incubated 20min at room temperature. The nanoparticles were washed again and resuspended in DropMap buffer containing fluorescent reporter proteins in a final concentration of 1.25 mg/ml beads. Reporter proteins were: Alexa647-labeled F(ab’)_2_ fragment of rabbit anti-human IgG Fc-specific (Jackson ImmunoResearch) used at 75nM final in-droplet concentration; Alexa488-labeled (ThermoFisher) GPIIbIIIa (purified protein, Enzyme Research) used at 30 nM final in-droplet concentration.

### Droplet production and collection

Droplets were generated using hydrodynamic flow-focusing on a microfluidic chip as described^2^. The wafer master of SU-8 photoresist layer (MicroChem) with ~40 μm of thickness were manufactured using soft-lithography^3^ and microfluidic chips were fabricated using poly-dimethylsiloxane (PDMS; Sylgard)^2^. Microfluidic chips were manufactured using soft-lithography in poly-dimethylsiloxane (PDMS). The continuous phase comprised 2% (wt/wt) “008 Fluorosurfactant” (RAN Biotechnologies) in Novec HFE7500 fluorinated oil (3M). Aqueous phases I and II were co-flowed and partitioned into droplets. The flow rate of aqueous phases I and II was 70 μL/h, whereas that of oil was 600 μL/h in order to achieve monodisperse droplets of ~40 pL volume. Newly generated droplets were directly injected into the DropMap 2D chamber system^2^, mounted on a fluorescence microscope (Ti Eclipse, Nikon). The emulsion was exposed to a magnetic field, forcing the nanoparticles inside each droplet to form an elongated aggregate termed “beadline”.

### Data acquisition

Images were acquired using a Nikon inverted microscope with a motorized stage (Ti-Eclipse). Excitation light was provided by a light-emitting diode (LED) source (SOLA light source, Lumencor Inc.). Fluorescence for the specific channels were recorded using appropriate band-pass filters and camera settings (Orca Flash 4.0, Hamamatsu) at room temperature and ambient oxygen concentration. Images were acquired using a 10x objective (NA 0.45). An array of 10×10 images was acquired for each replicate, every 7.5 min in all channels over 37.5 min (6 measurements total). Duplicates or triplicates were systematically acquired for every sample, with each replicate being the filling of the DropMap 2D chamber with a novel droplet population acquired over time on a 10×10 image array.

### Image analysis and calculations

Images were analyzed using a custom-made Matlab script (Mathworks) that identifies each droplet and the beadline within each droplet. Fluorescence values associated to the beadline is extracted as well as the mean fluorescence of the entire droplet except the beadline (background fluorescence). A value of fluorescent relocation for each droplet at each timepoint was calculated i.e., fluorescence value of the beadline divided by the fluorescent value of the background. Data was exported to Excel (Microsoft) and sorted for droplets showing an increase in relocation of the anti-IgG reporter fluorescence (Alexa647) over time and above a threshold of Alexa647-relocation > 1.5. The sorted droplets were visually controlled for the presence of a single cell within the droplet, for droplet movement between image acquisitions, absence of fluorescent particles other than the beadline (e.g., Alexa647-anti-IgG or Alexa488-GPIIbIIIa protein aggregates, cell debris) and undesired aggregation of fluorescent reporters on the cell surface inside the droplet. IgG secretion rate and dissociation constant (K_D_) were estimated as described^2^.

*Estimation of IgG concentration within each droplet*: a calibration curve for the estimation of IgG concentration was obtained by preparing droplet populations containing all bioassay reagents except cells that were replaced by a range of concentrations of monoclonal IgG1 (one concentration per droplet population). Images were acquired exactly 3min the droplets were immobilized in the DropMap chamber for each IgG concentration, and fluorescent relocation calculated. The calibration curve was subsequently used to estimate the IgG concentration and secretion rate (IgG molecules per second) for each time interval, and the mean secretion rate was calculated by averaging the values of all intervals.

*Estimation of dissociation constant (K_D_) for GPIIbIIIa within each droplet*: a calibration curve was obtained by defining relocation values for both anti-IgG (AF647) and GPIIbIIIa (AF488) for a collection of 10 anti-GPIIbIIIa mAbs with a range of K_D_ over 1 log as defined using Bio-layer interferometry (ForteBio). Droplet populations were generated for different concentrations of each mAb to be analyzed by the DropMap bioassay. A curve was defined by plotting the relocation from the anti-IgG (AF647) against relocation of GPIIbIIIa (AF488), and the slope of the resulting line was calculated and termed “DropMap slope”. The calibration curve was defined by the linear relation between K_D_ for GPIIbIIIa and the DropMap slope of each mAb, and allowed to extract a K_D_ value for each DropMap slope value calculated from a droplet containing an IgG-SC of unknown affinity for GPIIbIIIa.

### Affinity determination of monoclonal antibodies

Bio-layer interferometry measurements were performed using anti-human IgG sensors in an Octet system (ForteBio). Anti-GPIIbIIIa mAbs (10μg/mL) were captured on the sensors for 10 min. Equilibrium dissociation constants (K_D_) were determined by monitoring over 85 min the association between the immobilized antibodies and GPIIbIIIa in solution in duplicate for seven concentrations of antigen (200, 100, 50, 25, 12.5, 6.25 and 3.13 nM). An irrelevant IgG monoclonal antibody was used as negative control to subtract the background signal. Data analysis was performed using the Octet Analysis software (ForteBio), and the K_D_ were calculated by steady state analysis.

### Flow cytometry

Fresh PBMCs were isolated from venous blood samples via standard density gradient centrifugation. For the surface stain, cells were washed and resuspended at 2×10^6^ in 100μl PBS with 2% FBS and incubated with Zombie violet fixable viability dye (Biolegend) and an antibody cocktail for 25min at 4°C in the dark. Following surface staining, cells were fixed/permeabilized for 30min at 4°C in the dark with eBioscience FoxP3 transcription factor buffer kit and incubated with antibodies recognizing intracellular targets for 30min at 4°C in the dark. Samples were acquired on a LSR Fortessa (BD Biosciences). Data were analyzed with Kaluza software (Beckman Coulter). A list of antibodies used in this panel can be found in Table S1 and detailed gating strategy is depicted in Figure S5.

### ELISPOT

Anti-GPIIbIIIa ELISPOTS were performed as previously described^4^. Briefly, keyhole limpet hemocyanin (KLH, 2.5 μg/ml), goat anti-human Ig polyvalent antibody (10 μg/ml; Invitrogen), or purified GPIIbIIIa (15 μg/ml; Stago) were coated in phosphate-buffered saline (PBS) and 0.05% CaCl2 in multiscreen 96-well filter plates (MSIPS4510, Millipore) with overnight incubation at 4°C.

Then, 10^6^ splenocytes were serially diluted in culture medium (RPMI-1640 (Invitrogen) supplemented with 10% HyClone FBS (Thermo Scientific) and 1% Pen/Strep (Thermo Fisher)) in triplicate before transferring to ELISPOT plates and incubated overnight at 37°C with 5% CO2. Cells were removed, and the ELISPOT plate was incubated for 4 hours at 4°C with biotinylated goat anti-human IgG Fc (Invitrogen), followed by incubation for 1 hour at room temperature with horseradish peroxidase (HRP)–conjugated avidin (Vector Laboratories). HRP activity was further revealed using 3-amino-9-ethylcarbazole (BD Biosciences) for 8 min at room temperature in the dark. Spots were enumerated in each well with an ELISPOT reader using the AID software (version 3.5; AutoImmun Diagnostika).

### Statistical analyses

The R environment v4.0.5 was used for all the analyses^5^. Data were neither averaged nor normalized prior analyses. Response variables were log2 converted for better adjustment to linear models. Data were fitted to a linear (quantitative response) or logistic (qualitative response) model that includes the variables of interest, i.e., Group (HD or ITP classes) and Sample Type (SP, BL or BM classes), plus the Patient variable nested into the Group variable. Group and Sample Type interaction was added in the model related to Figure 2a because of the post-hoc inter-variable contrast comparisons. Otherwise, it was removed from the models when the effect was not significant. Age and Sex covariates were not incorporated as their effects were already represented by the Patient variable. Mixed models were used using the lmer() function of the lme4 package in order to consider the effect of Patient and patient:Group interaction as random. Anova analyses were performed with the Anova() function of the car package. Type 3 Sum of Squares was applied because of unbalanced designs. Two by two effect comparisons (contrast comparisons) were performed with the emmeans() function of the emmeans package. Unequal variance t test (Welch test) was used in bivariate designs, i.e., when data were analysed on a single patient (Figure 1) or in the presence of a single value per patient (Figure 3a). In figure 3d-f, a Pearson correlation test was performed after removal of zero values, log2 transformation, and residuals checking of the linear regression carried out in both ways. Statistical significance was set to P ≤ 0.05. In each figure, type I error was controlled by correcting the p values according to the Benjamini & Hochberg method (“BH” option in the p.adjust() function of R). Results are detailed in Table S2.

